# Hsp70/Hsp90 organizing protein (HOP) maintains CRAF kinase activity and regulates MAPK signaling by enhancing Hsp90-CRAF association

**DOI:** 10.1101/2023.02.17.528950

**Authors:** Nilanjan Gayen, Sahana Mitra, Somesh Roy, Atin K. Mandal

## Abstract

The stability and activity of CRAF kinase are stringently regulated by heat shock protein 90 (Hsp90). Hsp90-mediated client folding and maturation is governed by its co-chaperones, but their functionality in chaperoning CRAF/Raf1 kinase to accomplish signaling under physiological conditions remains poorly understood. Here, we show that Hsp70/Hsp90 organizing protein (HOP) associates with CRAF kinase for maintaining its kinase activity and facilitates the activation of the MAPK pathway. Such activation is mediated by TPR2A-2B-DP2 domain of HOP and requires efficient binding to Hsp90. Being a recruiter of Hsp90, Cdc37 is unable to supplement the function of HOP/Sti1. Downregulation of HOP/Sti1 in yeast and *in vitro* cell culture significantly reduces the CRAF signaling. Our data suggest that Hsp90 is recruited to CRAF in two steps, separately initiated by co-chaperones HOP and Cdc37 respectively during CRAF folding/maturation, and again upon CRAF activation mediated by HOP during MAPK signaling. Therefore, HOP is a regulator of CRAF kinase during activation of MAPK pathway and serves as a sensor of growth signaling beyond its client folding and maturation function.

## Introduction

The three-tiered RAF-MEK-ERK signaling cascade is involved in cell differentiation, proliferation, survival, and oncogenic transformation. RAF kinases are the key mediators to relay the signal from plasma membrane to the nucleus (*1*). Among the three RAF isoforms, CRAF/Raf-1 is the most ubiquitously expressed isoform in individual tissue (*2*). CRAF is found to be essential for normal cell growth, development, and embryogenesis (*3, 4*). Being an essential connector between Ras and MEK in intracellular signaling, CRAF activity needs to be precisely controlled. Deregulation of CRAF activity leads to several maladies including rasopathies and cancer (*5, 6*).

CRAF activity is majorly regulated by protein-protein interaction, and phosphorylation-dephosphorylation events (*7, 8*). Phosphorylation at both Ser-259 and Ser-621 (the binding sites of scaffold protein 14-3-3) residues keeps CRAF as an inactive kinase at the cytosol (*9–11*). Upon mitogenic stimulation dephosphorylation of Ser-259 by protein phosphatase 2A activates CRAF kinase (*12*). Unlike BRAF, CRAF is additionally assisted by the molecular chaperone Hsp90 (*13, 14*). Therefore, perturbation of Hsp90 functioning significantly reduces CRAF signaling by promoting kinase degradation (*15, 16*). Hsp90 stabilizes CRAF by promoting its autophosphorylation at Ser-621, and recruitment of Hsp90 is essential for CRAF activation during MAPK signaling (*17*).

Hsp90 participates in the folding, stability, and maturation of a diverse set of “Client proteins” including kinases, receptors, and transcription factors (*18–21*). Therefore, the involvement of Hsp90 in several physiological processes makes this chaperone a potential therapeutic target (*22*). The function of Hsp90 is ATP-dependent and stringently regulated by its different co-chaperones, which impart functional directionality to Hsp90 (*23–26*). Therefore, a set of co-chaperones required during Hsp90-dependent client protein folding. Kinase chaperoning is principally governed by co-chaperone Cdc37 (*21, 27, 28*), while that of Glucocorticoid receptors (GR) is primarily regulated by Sti1/HOP (*29, 30*). However, yeast MAP3K kinase, Ste11 activity is regulated by both Sti1 and Cdc37 (*31*). In this context, Sti1/HOP acts as an ‘adaptor’ to coordinate the efficient client transfer from Hsp70 to Hsp90 by its tetratricopeptide repeat (TPR) motif (*32*). Moreover, HOP/Sti1 has a selective preference to interact with the apo form of Hsp90 during client loading step (*33*). Therefore, HOP interacts with Hsp90 client proteins and regulates their folding and activity. HOP clients are responsible for maintaining cell proliferation and notably, HOP upregulation is found in various cancer tissues (*34–36*). Previous findings establish that extracellular HOP regulates Hsp90α mediated MMP2 activation during breast cancer cell migration and invasion (*37*). However, the underlying mechanism of how intracellular HOP regulates CRAF activation during signaling remains unknown. Here, we show that HOP/Sti1 regulates CRAF basal activity and precisely controls signal-dependent CRAF kinase activation. We also observed that HOP exploits its Hsp90 interacting domain to regulate the CRAF functioning. Furthermore, our data suggests that CRAF chaperoning by HOP/Sti1 shares a non-redundant function with Cdc37. Therefore, this study signifies a novel molecular mechanism of intracellular HOP by which this co-chaperone functions as an essential component of CRAF signalosome to regulate the CRAF kinase activation during MAPK signaling.

## Result

### Sti1/HOP is essential for CRAF functioning

CRAF kinase stabilization and maturation are dependent on its c-terminus Ser-621 auto-phosphorylation (*17, 38*). Our previous study has shown that CRAF auto-phosphorylation is governed by both Hsp90 and Cdc37 (*17*). Hsp90 mediated client maturation and activation requires assistance from different co-chaperones including Aha1, Sti1, and Sba1 (*23–26, 39*) respectively. So far, Sba1 is reported to control the receptor activation (*40*), while role of Aha1 is established on several kinases including CRAF activation (*14*). Although Sti1/HOP is an essential component in Hsp90-mediated chaperone cycle, the potential contribution of Sti1/HOP in CRAF kinase functioning is not clear. To begin with, initially, the role of yeast Sti1 on human CRAF kinase maturation and function was investigated in *Saccharomyces cerevisiae*, since HOP and Sti1 function similarly (*41*). The cDNA encoding CRAF kinase was expressed under GPD promoter in different yeast co-chaperone knockout strains (as indicated in Fig. 1Ai). We observed that *sti1* deletion significantly abolished the basal kinase activity of the expressed CRAF kinase (Fig. 1Ai), while showing no effect on phosphorylation at Ser-621 or Ser-259 in CRAF kinase (Fig. 1Aii). However, *sba1* deletion did not show any significant effect on CRAF activity keeping the notion that Sba1 specifically controls the client receptor functioning. These data together suggest that yeast Sti1 might regulate the function of CRAF client independent of the kinase maturation event.

**FIGURE 1:**
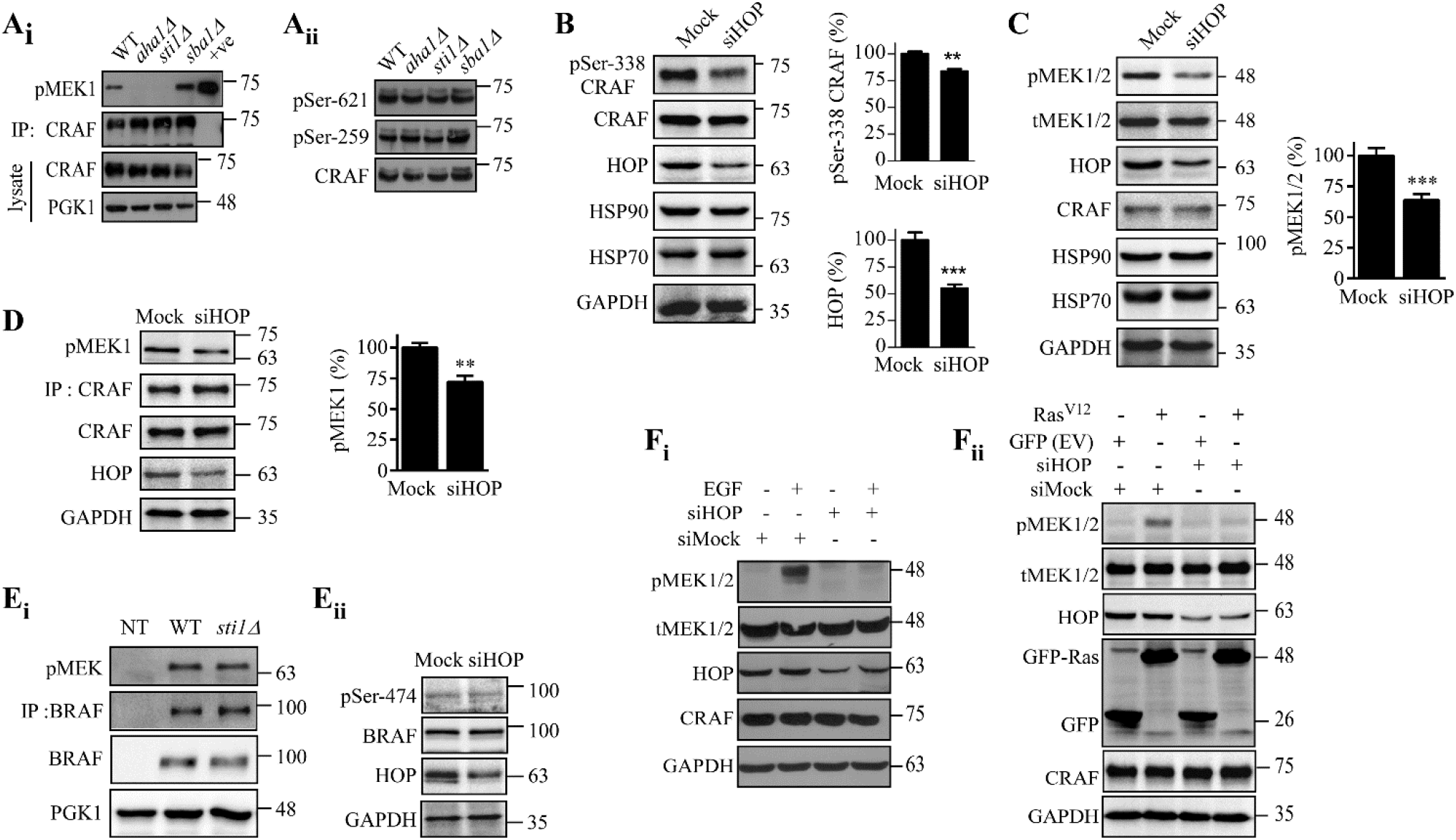
Sti1/HOP maintains CRAF kinase activity. **(A)**. Analysis of CRAF kinase activity and its basal phosphorylation from Hsp90-dependent co-chaperone knockout yeast strains. Flag-CRAF cloned under GPD promoter was expressed into yeast knock-out strains, as indicated. CRAF was pulled using anti-Flag antibody and *in-vitro* CRAF kinase activity was measured as described in the experimental procedures. Purified CRAF was used as positive (+ve) control of kinase assay. PGK1 was used as loading control (Ai). CRAF basal phosphorylation (pSer-621 and pSer-259) was monitored using phospho-specific antibodies upon loading equal amount of protein from the indicated co-chaperone knockout yeast strains (Aii). **(B-D)**. HOP knock down was performed by treating HEK293T cells with 100 nmoles of smart pool HOP siRNA (Dharmacon) for 96 hours. pSer-338 CRAF and pMEK1/2 levels were analyzed from the total cell lysates, quantified and plotted upon normalization by total CRAF (B) and total MEK1/2 (C). Endogenous CRAF was pulled and the basal kinase activity in HOP silenced condition was assessed by MEK phosphorylation, normalized by pulled CRAF protein and shown in the graph (D). Statistical analysis was determined by unpaired t-test, where ** and *** represent p<0.01 and p<0.001 respectively. **(E)** Sti1/HOP does not affect BRAF activity. p416-GDP-Flag-BRAF was expressed in *sti1Δ* yeast strain and *in-vitro* BRAF kinase activity was analyzed by MEK phosphorylation. Here, PGK1 represented the loading control. NT indicated non-transformed yeast cell (Ei). BRAF phosphorylation (Ser-474) was analyzed from HOP silenced HEK293T cells (Eii). **(F)** HOP silencing affects activation of MAPK pathway. HOP was silenced in HEK293T cells as described in (B-D). Cells were starved at last 24 h post-siRNA treatment, followed by the addition of 20ng/ml of EGF for 15 mins (Fi). Similarly, GFP-Ras^V12^ was transfected to HOP silenced cells at 48h post-siRNA treatment (Fii). At 96 h of siRNA transfection, cells were lysed and analyzed with respective antibodies (Fi and Fii). GAPDH was used as loading control in (B, C, D, Eii and F respectively). Results are representative of three independent experiments.

To confirm the analogy between the role of yeast Sti1 and its mammalian homolog HOP in CRAF functioning, further experiments were performed using *in-vitro* cell culture system. HOP silencing in HEK293T cells reduced CRAF activation-related phosphorylation at Ser-338 site, MEK phosphorylation, and *in-vitro* kinase activity respectively (Fig.1B-D). Similar observation was obtained for the hyperactive CRAF mutant S257L (Fig. S1) suggesting that the physical association of HOP with CRAF is essential to maintain the optimum functioning of the CRAF complex. In contrast, BRAF activity remained unchanged in Sti1/HOP depleted condition (Fig. 1Ei and Eii). Additionally, HOP silencing also reduced MEK activation either in the presence of potential activator of MAPK pathway, EGF (Fig. 1Fi) or Ras^G12V^ (Ras^V12^) (Fig. 1Fii). These data collectively demonstrate that co-chaperone HOP is required to maintain CRAF functioning during MAPK signaling.

### HOP over-expression activates RAF signaling through CRAF kinase

Previous studies have shown that the upregulation of certain Hsp90 co-chaperones such as Cdc37 and Aha1 activates RAF signaling pathway (*13, 14*). In this study, we explored the effect of time-dependent and dose-dependent HOP over-expression on MAPK signaling in HEK293T cells. In both conditions, MEK phosphorylation was increased (Fig. 2Ai and Aii). HOP over-expression also showed a cumulative activating effect either in presence of GFP-Ras^V12^ or EGF treatment (Fig. 2Bi and Bii) suggesting its potential role during MAPK signaling. Furthermore, extracellular HOP can also activate MAPK signaling in an endocytosis-dependent way (*42*). We checked the presence of any secreted fraction of HOP during our study by culturing the cells in serum-free conditioned media (CM) that facilitates the secretion of intracellular HOP (*43*). Absence of HOP in conditioned media (CM) ruled out the possibility of MAPK activation from secretory HOP and confirmed the CRAF activation during our study is due to HOP over-expression inside the cell (Fig. 2C). MAPK signaling is activated either by RAF kinase (*44*) or by PI3K kinase (*45*). To confirm the involvement of MEK as a mediator during HOP-induced MAPK activation, cells were treated with MEK inhibitor, U0126 (*46*). MEK inhibition suppressed the activation of MAPK pathway under HOP over-expressed condition (Fig. 2D). In parallel, being an Hsp90 client, MEK could also be directly activated by co-chaperone HOP. To exclude this possibility, cells were treated with RAF inhibitor sorafenib (*47*). Such treatment showed reduction in MEK phosphorylation suggesting that HOP-mediated MEK activation is not a direct effect, but involves RAF-MEK axis (Fig. 2E). Cumulatively, the observed data establishes that HOP has a contributory role during RAF signaling.

**FIGURE 2:**
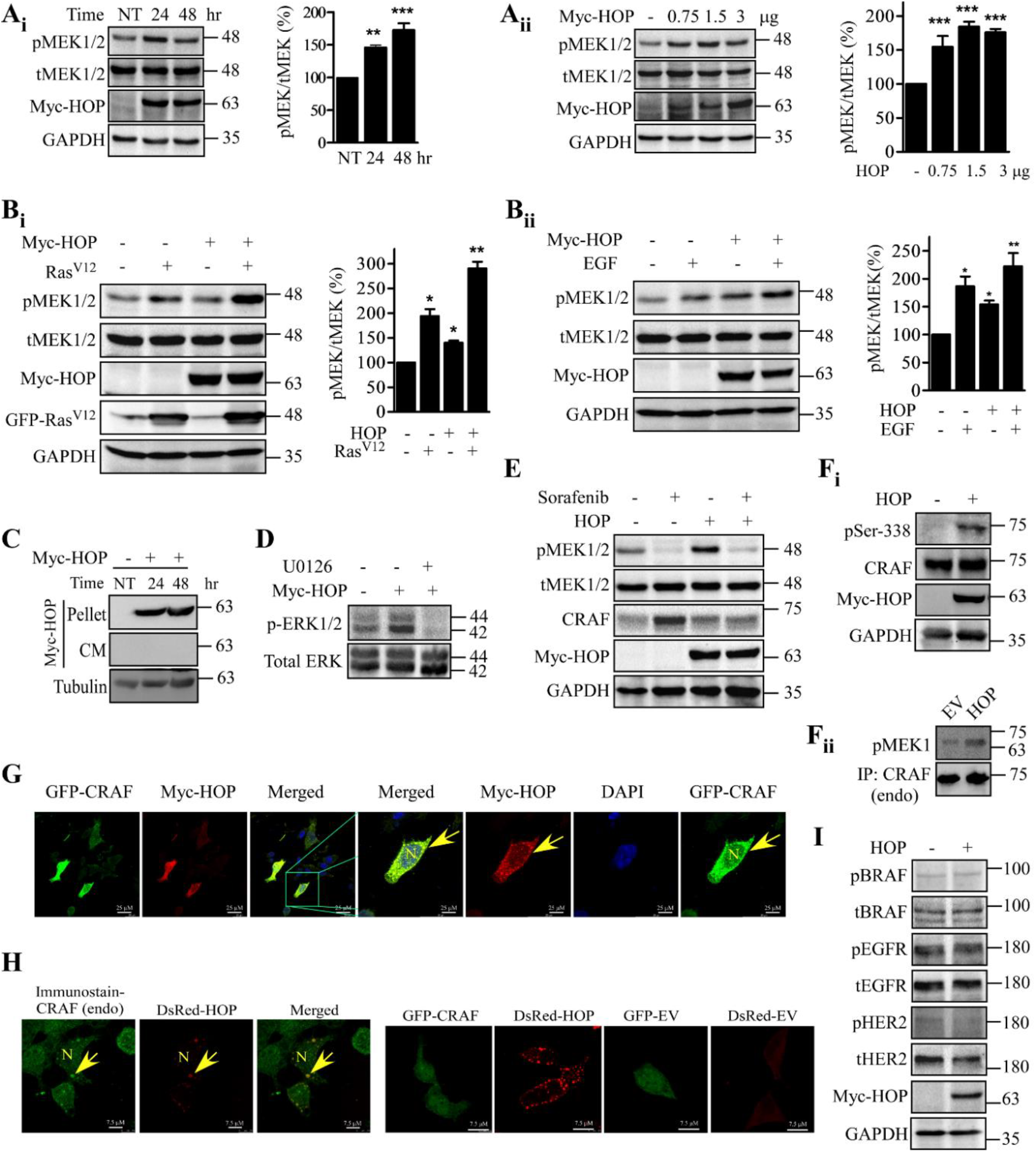
HOP enhances MAPK signaling by activating CRAF kinase. **(A)**. Time-dependent (Ai) and dose-dependent (Aii) Myc-HOP over-expression was performed in HEK293T cells. MEK activation was monitored using pMEK1/2 antibody. **(B)** HEK293T cells were transfected with Myc-HOP for 48 h either with mEGFP-Ras^V12^ (Bi) or treated with EGF as described before (Bii). Cell lysates were analyzed for MEK phosphorylation. Adjacent graph shows the quantification of pMEK1/2 normalized with total MEK1/2. Statistical analysis was performed by using one-way anova, where *, ** and *** indicate p<0.05, p<0.01 and p<0.001respectively. **(C)** Detection of the presence of over-expressed Myc-HOP in secreted fraction. HEK293T cells were transfected with Myc-HOP for 24 h and 48 h respectively. The culture media was immediately replaced by serum-free media. Conditioned media/ cultured serum-free media (indicated as CM) and cell pellets were then collected at 24h and 48h post-transfection. Secreted Myc-HOP was detected as described in the experimental procedures using anti-Myc antibody. Here, tubulin represented the loading control. **(D-E)** HEK293T cells transfected with Myc-HOP were treated with MEK inhibitor U0126 and RAF inhibitor sorafenib (5 μM, 1 hr) at 48h post-transfection. Cell lysates were checked for ERK and MEK phosphorylation respectively. **(F)** The CRAF activation was detected in Myc-HOP over-expressed HEK293T cells by analyzing the level of pSer-338 CRAF (Fi) and CRAF basal kinase activity from the same lysate by immunoprecipitating endogenous CRAF (Fii). **(G-H)** HEK293T cells were transfected with either GFP-CRAF and Myc-HOP together or DsRed-HOP alone for 48 hours. Localization of CRAF (GFP fluorescence) and HOP (immunostaining with anti-myc antibody) was analyzed under confocal microscope (G). Endogenous CRAF was immunostained with anti-CRAF antibody upon over-expression of DsRed-HOP in HEK293T cells. Confocal images of the empty vectors (GFP-EV and DsRed-EV respectively), GFP-CRAF, and DsRed-HOP were shown as control (H). Representative fields are shown. Yellow arrowhead represented the CRAF clusters that are confined to the close proximity of plasma membrane in HOP over-expressed cells. **(I)**. WB analysis showed the activation status of BRAF, EGFR and HER2 with phospho specific antibodies in HOP over-expressed HEK293T cells. GAPDH was used as loading control. Results were obtained from three independent experiments.

RAF kinases are the key mediators to phosphorylate and activate MEK (*44*). So far, our data suggested that HOP particularly controls CRAF functioning. Therefore, we checked CRAF activation in presence of exogenously expressed HOP in HEK293T cells. The involvement of CRAF kinase in HOP-mediated MAPK activation was confirmed by the increased Ser-338 phosphorylation and the basal kinase activity respectively (Fig. 2Fi and Fii). Furthermore, confocal data showed a significant change in the subcellular localization of GFP tagged CRAF kinase upon Myc-HOP over-expression. Increased localization of GFP-CRAF at the cell periphery indicates the activation of CRAF kinase (Fig. 2G), as shown by the previous studies also (*48*). In consistent with this data, we also observed that the endogenous CRAF localizes at the cell periphery upon DsRed-HOP over-expression (Fig. 2H). Earlier studies have shown that growth signal is transmitted through CRAF/BRAF dimer (*49*). Therefore, to exclude the possible involvement of BRAF kinase during HOP-induced MAPK signaling, BRAF activation was assessed by WB analysis. The experiment showed that neither BRAF nor its upstream kinase, either EGFR or HER2 is activated by HOP over-expression (Fig. 2I). These findings collectively demonstrated that HOP is essential for efficient MAPK pathway activation *via* CRAF kinase.

### TRP2A-2B-DP2 domain of HOP ensures Hsp90 mediated CRAF activation

HOP functioning is regulated by several phosphorylation events (*41*). The phosphorylation sites are present either in or at the flanked region of Hsp70/Hsp90 interacting domains, primarily TPR2A and TPR2B, or in the linker region connecting the main modules (Fig. 3A) (*50*). Previously, it has been shown that the mutation at Y354, T198, and K229 of mammalian HOP reduce the interaction with Hsp70 and Hsp90 respectively (*51–53*). Our co-IP data is also in line with these previous observations (Fig. S2). Individual over-expression of either T198E or K229A mutants showed reduction in MEK activation in comparison to either WT or Y354E mutant in HEK293T cells (Fig. 3B). Similar to WB analysis for MEK activation in presence of HOP mutants as indicated, confocal imaging further showed a significant reduction in CRAF cluster at the cell periphery in case of T198E, and K229A alone and the double mutant (T198E, K229A) respectively. On contrary, Y354E expressing HEK293T cells showed a slight reduction in the CRAF cluster formation in comparison to wild type (WT) control (Fig. 3C and Fig. S3). Similar result in reduction of p-Tyr activity was observed for Y354E mutant (*53*). Furthermore, the co-IP analysis from both cell culture and yeast showed that the mutation at T198 and K229 position of HOP abrogated the physical interaction between HOP and CRAF (Fig. 3D and Fig. S4A-B). Altogether, these observations suggested that the interaction between HOP and Hsp90 is required for efficient CRAF activation during MAPK signaling.

**FIGURE 3:**
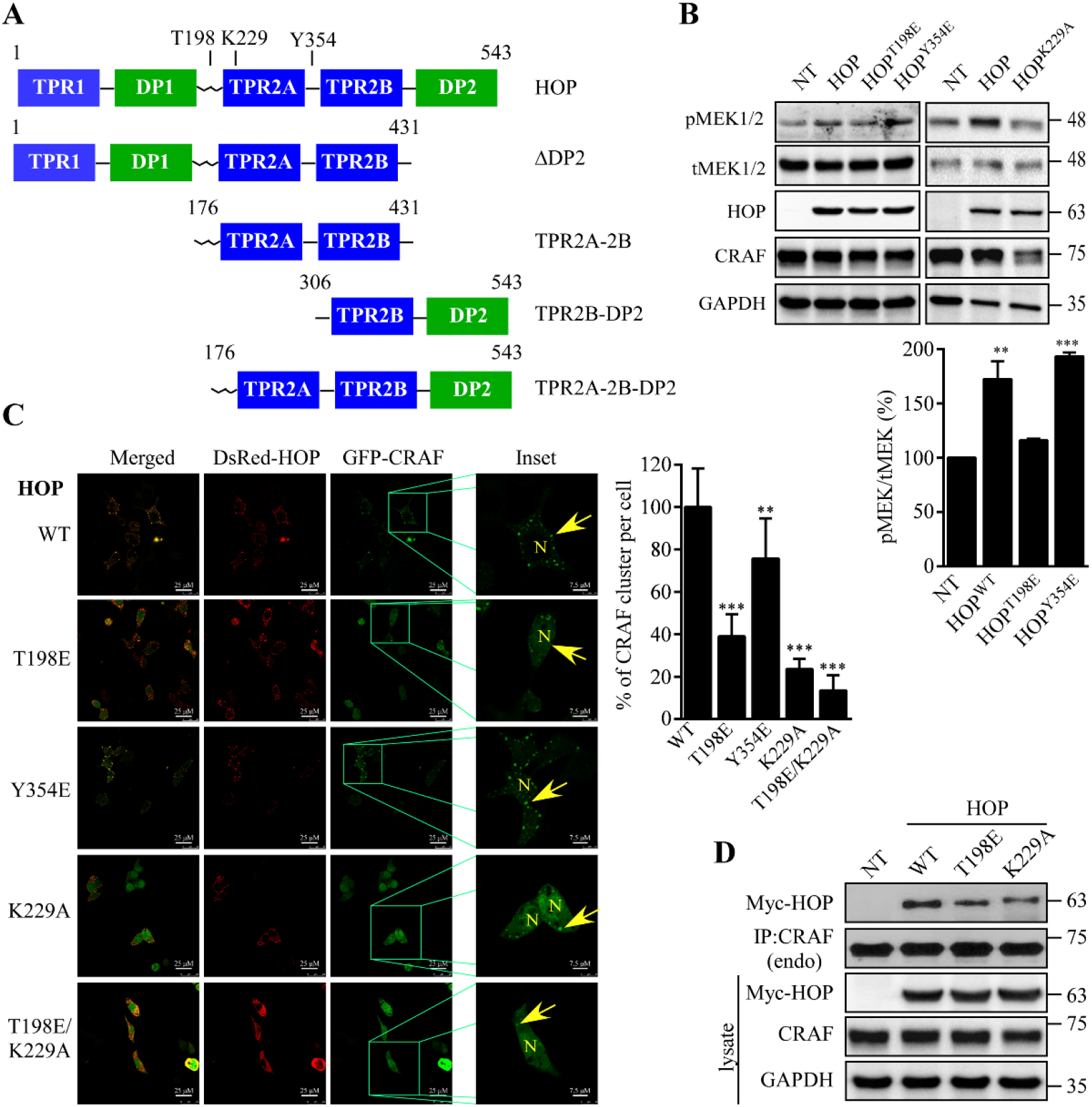
Hsp90-HOP interaction is crucial for CRAF activation. **(A)** Domain organization of HOP highlighting the residues, T198, K229 and Y354 responsible for Hsp90 and Hsp70 binding respectively. **(B)** HEK293T cells transfected with either Myc-tagged WT or other HOP mutants (indicated in figures) were lysed, analyzed by WB to monitor MEK phosphorylation using pMEK1/2 antibody and plotted by normalizing with total MEK1/2 as shown in the adjacent graph. HEK293T transfected with GFP-CRAF and DsRed tagged either WT or mutant HOP were imaged to identify the distribution of CRAF cluster. Scale bar 25 μm. Yellow arrowhead in the inset (~3.5X magnificationof a single cell with scale bar 7.5 μm) indicates the formation of CRAF cluster close to the membrane in the respective fields. The adjacent graph represents quantification of the percentage of CRAF cluster per cell, mean of clusters quantified from ≥10 cells from three independent experiments. Statistical analysis was performed by one-way anova, where ** and *** indicate p<0.01 and p<0.001 respectively. **(D)** Endogenous CRAF was pulled from HEK293T cells transfected with either Myc tagged WT or mutant HOP and blotted using anti-Myc antibody. GAPDH was used as loading control. NT indicates non-transfected HEK293T cell.

HOP comprises three TPR domains, organized in two modules containing one and two TPR domains followed by a DP domain (Fig. 3A) (*50*). The M domain of Hsp90 interacts with TPR2A, while TPR1 and TPR2B are responsible to interact with Hsp70 and DP domain contributes to *in-vivo* client activation (*54, 55*). The interaction between TPR2A and DP2 domain is essential for *in-vivo* client activation shown previously for the GR receptor (*41*). Our data from both yeast and *in-vitro* cell culture systems suggests that TPR2A-2B module by itself is unable to support CRAF activation. Similarly, deletion of either TPR2A or DP2 from mammalian HOP showed reduction in CRAF activation. However, the presence of both modules TPR2A and DP2 along with TPR2B significantly increased CRAF activity (Fig. 4A and B) indicating that TPR2A-TPR2B-DP2 fragment is indispensable to maintain CRAF activation. The TPR2A-2B-DP2-mediated activation of CRAF kinase was substantially reduced when HOP-Hsp90 interaction was compromised by introducing K229A mutation (Fig. 4B). Furthermore, co-IP analysis suggests that HOP is an intrinsic component of CRAF complex (Fig. 4C and Fig. S5) and TPR2A-2B-DP2 region of HOP is essential to keep CRAF-Hsp90 association intact (Fig.4D). Interestingly, abolition of Hsp90 interaction with HOP^TPR2A-2B-DP2^ by insertion of T198E or K229A mutation significantly reduced CRAF-Hsp90 association. Notably, blocking of HOP^TPR2A-2B-DP2^-Hsp90 interaction using double mutant (T198E,K229A) leads to the complete abolition of CRAF-Hsp90 association (Fig.4Dii), while interference of HOP^TPR2A-2B-DP2^-Hsp70 interaction using HOP^TPR2A-2B-DP2^Y354E mutant has no significant contribution to reduce the residual CRAF-Hsp90 assemblage (Fig.4Di). It further establishes that Hsp70 is not involved to hold CRAF-Hsp90 complex. This observation is in parallel with the previous study (*53*) showing that the modification at Y354 of HOP is not essential for Hsp90-Hsp70 assemblage with the client. These data also indicate the fact that the recruitment of both HOP and Hsp90 to CRAF complex is a simultaneous process during CRAF activation. Altogether, the data suggests that HOP remains associated with CRAF complex in conjugation with Hsp90 in presence of stimulus during MAPK signalling.

**FIGURE 4:**
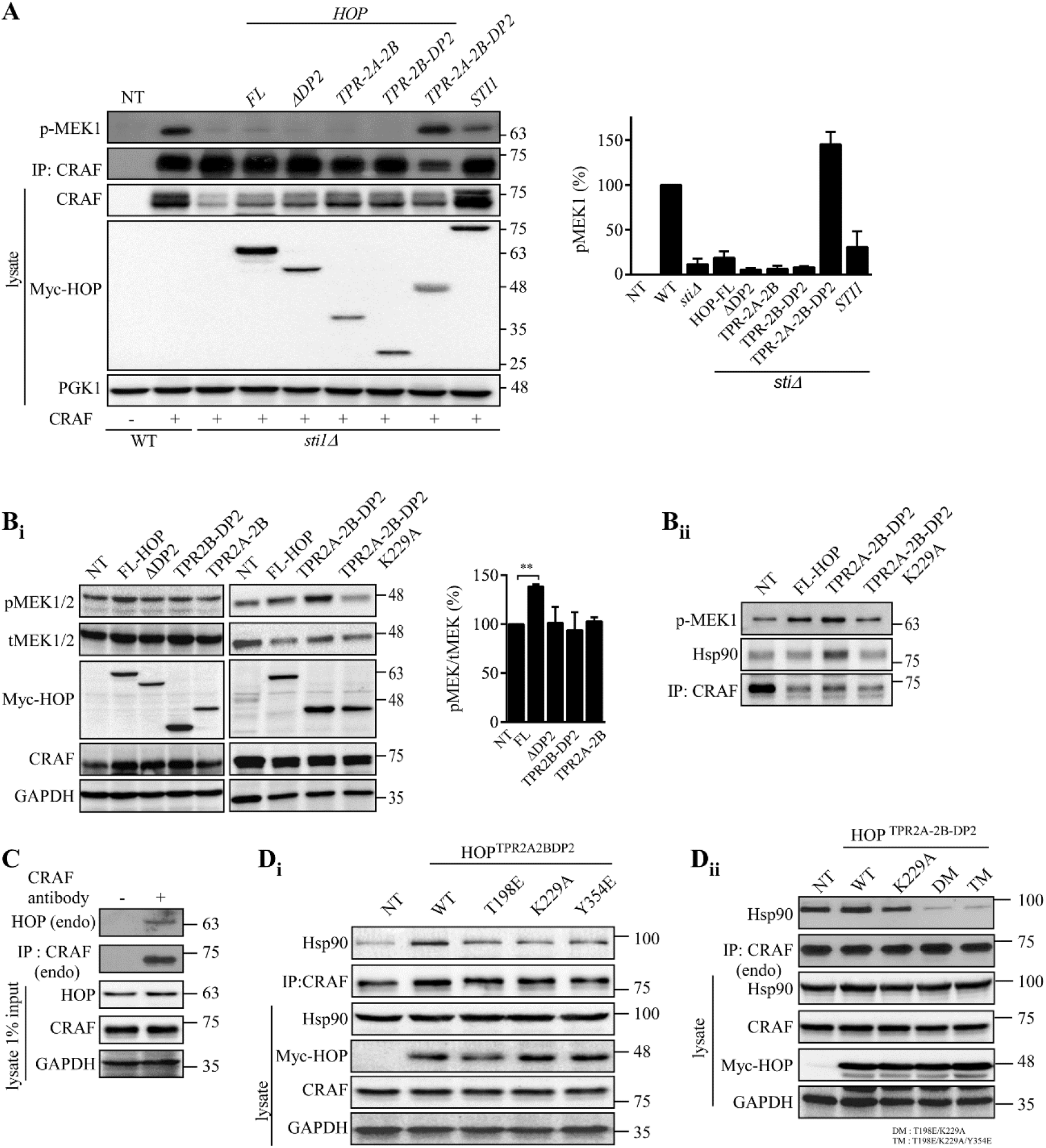
TRP2A-2B-DP2 domain is essential for Hsp90 mediated CRAF activation. **(A)** *sti1Δ* yeast cells were co-transformed with the indicated Myc tagged HOP/STI1 constructs and Flag-CRAF. CRAF was pulled using anti-CRAF antibody and *in-vitro* CRAF kinase assay was performed. Quantification of kinase activity was plotted as % of MEK1 phosphorylation normalized with the pulled CRAF, shown in the adjacent graph. PGK1 represented loading control. **(B)** MEK phosphorylation was checked in HEK293T cells upon transfection of Myc-HOP constructs as indicated in the figure. Adjacent graph represented the quantification of MEK1/2 phosphorylation normalized with total MEK1/2 (Bi). CRAF kinase activity was also checked (Bii). **(C)** Endogenous CRAF was pulled from HEK293T cell and HOP interaction was monitored by WB using anti-HOP antibody. **(D)** Association of CRAF with TPR2A-2B-DP2 domain of HOP and its corresponding mutants as indicated in the figure. Here, endo stands for endogenous protein. GAPDH was used as loading control. One-way anova was used for statistical analysis, where ** represent p<0.01. Similar results were obtained in three independent experiments except for K229A mutant. NT in Fig (B and D) indicates non-transfected HEK293T cell, while NT in Fig. A stands for non-transformed yeast cell.

### HOP and Cdc37 play a non-redundant role in CRAF kinase quality control

Selective entry of co-chaperones into the Hsp90 chaperone cycle regulates client quality control and regulation of signaling pathways. Hsp90 gains the substrate specificity by the recruiter co-chaperone Cdc37 (*21, 27, 28, 56*). Being a stringent client of Hsp90, post-translational maturation of CRAF kinase is dependent on both Hsp90 and Cdc37 (*17*). Our previous data established that the presence of Hsp90 in CRAF complex is maintained by Cdc37 (*17*). However, our present study suggests that another co-chaperone of Hsp90, HOP is also essential to maintain CRAF activation, and the interaction between Hsp90 and CRAF was reduced under HOP-depleted condition (Fig. 5A). These observations strengthen the fact that CRAF post-translational maturation (Ser-621 phosphorylation) is precisely dependent on co-chaperone Cdc37 (Fig. 5Bi and Bii), while HOP exhibits post-maturation function on CRAF kinase reflected from the CRAF *in-vitro* kinase assay (Fig. 1A and D). Overexpression of Cdc37 (mammalian/yeast) cannot supplement the loss of CRAF activity observed in *sti1* knockout yeast background (Fig. 5Ci and Cii), although over-expression of Cdc37 activates CRAF kinase as evident in Fig. 5D and from the previous study (*17*). Intriguingly, HOP over-expression also upregulates CRAF activation without influencing its Ser-621 phosphorylation (Fig. 5Bi and D). However, our present study shows that Cdc37 is unable to activate CRAF kinase under HOP-depleted condition (Fig. 5E). This observation suggests that Cdc37-mediated CRAF activation is governed by the degree of CRAF Ser-621 phosphorylation, while the presence of HOP in the CRAF complex is essential to keep the kinase in its active state. Therefore, these observations reflect a non-redundant behavior of Hsp90-related co-chaperones in client-specific context.

**FIGURE 5:**
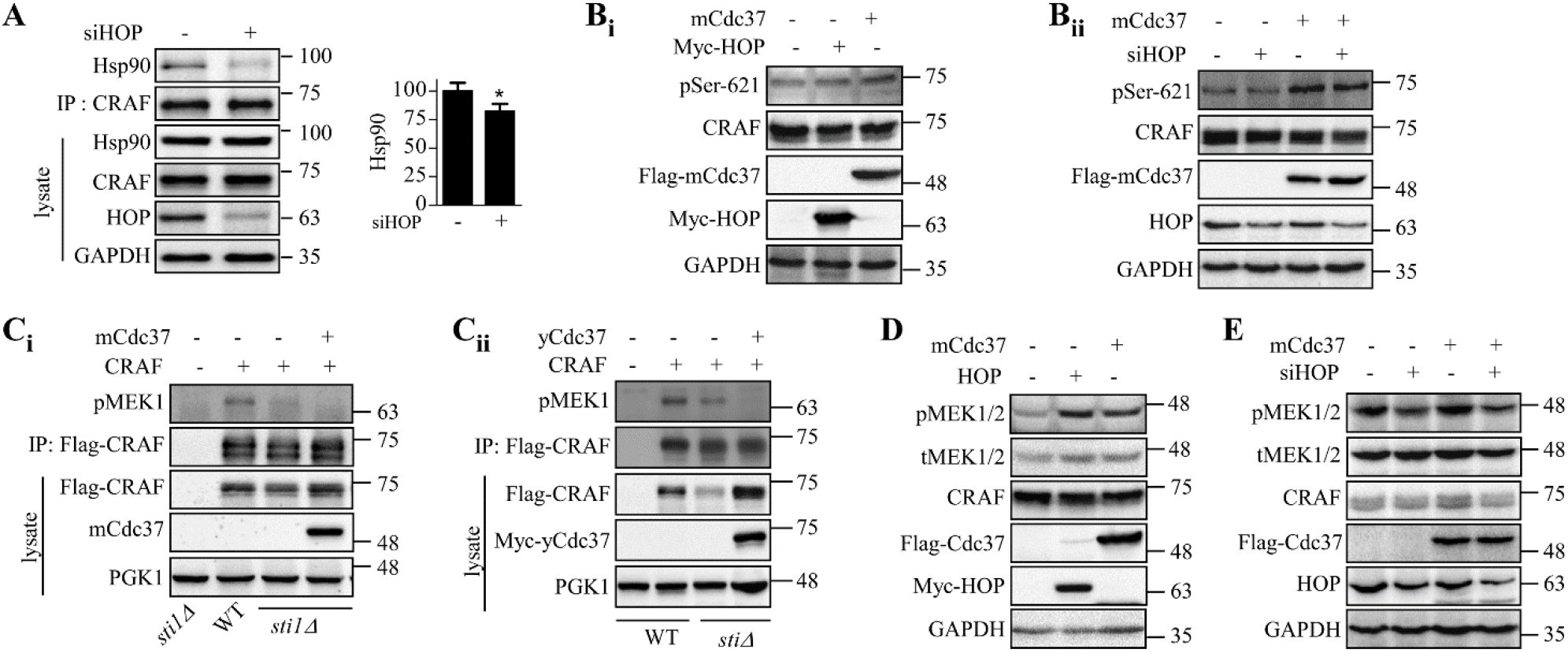
Non-redundant function of HOP and Cdc37 in CRAF kinase maturation and activation. **(A)** Immunoprecipitation was carried out by pulling down endogenous CRAF using anti-CRAF antibody and pulled samples were blotted to detect Hsp90 bound to CRAF. Anti-HOP antibody was used to check the level of HOP knock down in WB. **(B)** To determine the effect of HOP on CRAF Ser-621 phosphorylation (maturation marker), HEK293T cells were transfected with Myc-HOP and Flag tagged mammalian Cdc37 (mCdc37) respectively (Bi). Similar transfection reaction was performed under HOP silenced condition (Bii). At post transfection hour, cells were lysed and analyzed by WB to detect CRAF Ser-621 phosphorylation using pSer-621 antibody. (**C)** CRAF *in-vitro* kinase assay was performed by pulling exogenously expressed Flag-CRAF from *sti1Δ* yeast cells transformed with either Myc-mCdc37 (Ci) or Myc-yeast Cdc37 (Myc-yCdc37) (Cii). CRAF activity was assessed by monitoring MEK1 phosphorylation using anti-pMEK1 antibody in WB analysis. As an IP control, sti1 knockout lysate was taken. Here, PGK1 was taken as loading control. **(D-E)** MAPK pathway activation was analyzed from HEK293T cells transfected in a similar way, described previously (5B) using anti-pMEK1/2 antibody. In (A, B, D and E) GAPDH represented loading control. Similar results were obtained from three independent experiments.

### HOP engages both Hsp90 and actin in CRAF complex during MAPK signaling

In cancer cells, increased level of co-chaperones renders client maturation and activation (*34, 35, 57, 58*). Previous study has shown that the translocation of cytosolic CRAF during MAPK signaling is Hsp90-dependent and requires engagement of actin (*17*), here whether Hsp90 requires HOP as a co-chaperone for CRAF trafficking that needs to be examined. To test this, the association between Hsp90 and CRAF kinase was examined under differential expression of HOP. WB analysis revealed that gradual increment in cellular HOP level recruited Hsp90 to CRAF (Fig. 6A). Furthermore, impairing HOP-Hsp90 association by inserting point mutation in HOP either with T198E or K229A showed reduction in interaction between CRAF and Hsp90 (Fig. 6B). Interestingly, enhanced HOP recruitment to CRAF was observed upon activation of MAPK pathway by Ras^V12^ over-expression or EGF treatment (Fig.6C). In addition, HOP over-expression also revealed enhanced recruitment of actin and Hsp90 to CRAF complex (Fig. 6D), but not with protein translocator, HDAC6 (Fig. S6). Previously, it has been shown that both Hsp90 and actin are recruited to CRAF complex during its activation (*17*), but how co-chaperones assist Hsp90 to form this complex is still elusive. Our present data suggests that CRAF activation requires increased physical association with cytosolic Hsp90 and this interaction is HOP-dependent. In addition, reduction in Hsp90-CRAF association during Ras^V12^ -mediated MAPK signaling under HOP silenced condition further ensures the simultaneous recruitment of both HOP and Hsp90 during CRAF activation (Fig. 6E). Collectively, these results establish that CRAF signal complex requires both Hsp90 and HOP to maintain precise trafficking of the active CRAF by engaging cytoskeletal protein actin during MAPK signaling.

**FIGURE 6:**
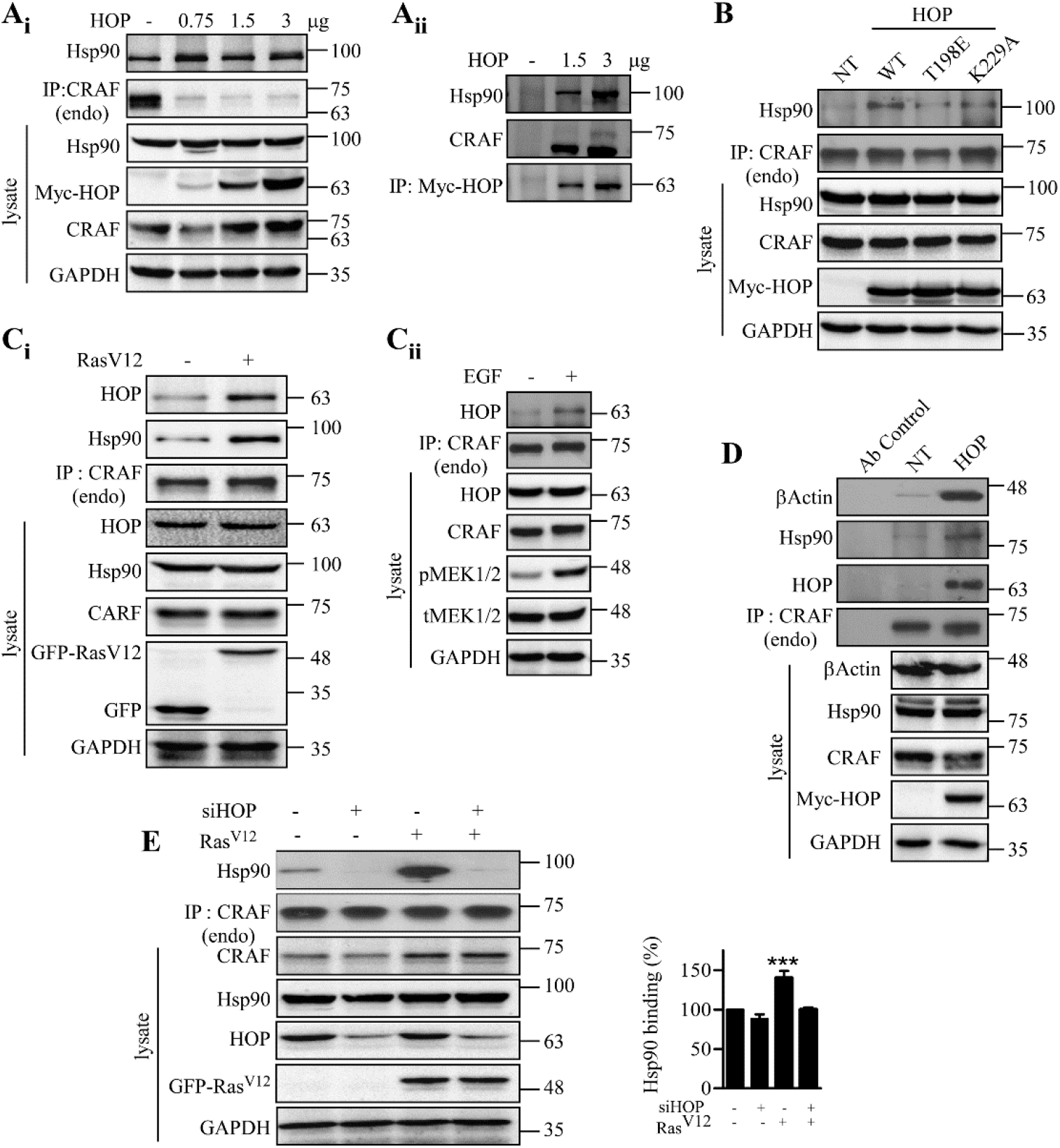
HOP facilitates interaction of CRAF with Hsp90 and actin during activation of MAPK signaling. **(A)** HEK293T cells were transfected with gradual over-expression of Myc-HOP. Cell lysates were immunoprecipitated by pulling either endogenous CRAF (Ai) or Myc-HOP (Aii). Protein interactions were identified by WB using anti-Hsp90 and anti-CRAF antibodies respectively (Ai and Aii). **(B)** Cells transfected with Myc-tagged HOP bearing indicated mutations T198E, and K229A, were lysed and endogenous CRAF was pulled using anti-CRAF antibody. IP samples were analyzed by WB to detect the level of bound Hsp90 using anti-Hsp90 antibody. **(C)** To activate RAF signaling, cells were either transfected with mEGFP-Ras^V12^(Ci) or serum starved followed by the EGF treatment (20 ng/ml) for 15 mins (Cii). After cell lysis, endo CRAF was immunoprecipitated and assessed to analyze CRAF-bound HOP using anti-HOP antibody. **(D)** Endogenous CRAF was pulled from HEK293T cells transfected with either empty vector (EV) or Myc-HOP. The IP samples were analyzed by WB to determine CRAF bound actin, Hsp90 and HOP using anti-β actin, -Hsp90 and -HOP antibodies respectively. **(E)** HEK293T cells were transfected with HOP siRNA or scrambled siRNA for 96 hours. mEGFP-Ras^V12^was over-expressed at 48 h post siRNA transfection. Endogenous CRAF was pulled and assessed for Hsp90 binding using anti-Hsp90 antibody. Here, GAPDH was used as loading control in cell lysate. Quantification of associated Hsp90 to CRAF is plotted in the adjacent graph. Significance was calculated by one-way anova. *** indicate p<0.001. Representative blots were from three independent experiments. NT, non-transfected control cell.

## Discussion

Aberrant regulation of Ras-RAF pathway can contribute to several abnormalities such as autocrine transformation, drug resistance, senescence, premature aging, and metastasis (*59*). Since, hetero-dimer of RAF isoforms predominantly transduce signal via MAPK pathway (*49*) therefore, controlling abnormal RAF regulation by using kinase-specific inhibitors becomes ineffective due to transactivation of other RAF isoforms (*60–62*). Although, BRAF is the most active isoform, however, CRAF is found to be the key mediator in transducing signal along MAPK pathway (*63, 64*). Therefore, ablation of CRAF level induces regression of *Kras/Trp53* driven tumors (*65–67*). Notably, kinase dead CRAF mutant is capable in pathway activation and causes disease progression (*68*). Thus, identifying the regulator of CRAF is crucial to modulate the MAPK signaling. So far, several regulators of CRAF have been identified that belong to different molecular categories including scaffold protein, kinase suppressor and molecular chaperones (*13, 69–71*). Among the chaperones, Hsp90 and Cdc37 are the well-characterized folding components of CRAF kinase (*13, 14*). Our previous study established that Hsp90 aids in CRAF folding by facilitating its Ser621 phosphorylation necessary for CRAF stability and activity. Additionally, recruitment of Hsp90 activates CRAF during signaling via MAPK pathway which reinforces to study the role of Hsp90 recruiter, the co-chaperones of Hsp90. Among the co-chaperones of Hsp90, HOP/Sti1 is identified as a regulator of different Hsp90 clients including steroid hormone receptors (SHRs), kinases v-Src, CHK1 and Ste11 in yeast (*31, 72–77*). Outside the context of protein folding, HOP/Sti1 is engaged in several biological processes including embryoid body formation, invasion of cancer cells, dicer independent loading of duplex RNA, growth, and survival of hepatic carcinoma cell (*37, 78–80*). However, the RAF kinase regulation by HOP/Sti1 in eukaryotes so far remains poorly defined.

The work presented here demonstrates that HOP/Sti1 acts as a recruiter of Hsp90 to the cytosolic CRAF complex during MAPK signaling. HOP/Sti1 as a co-chaperone assists the client transfer process from Hsp70 to Hsp90 during the ‘late complex’ formation in the protein folding cycle by keeping Hsp90 in an open conformation (*55, 81*). Therefore, perturbation of HOP/Sti1 functioning impairs the folding and maturation of Hsp90 client proteins e.g. Ste11 kinase, GR receptor, and Src kinase respectively indicating that HOP/Sti1 performs a significant contribution in the pre-maturation step during Hsp90-mediated client folding cycle (*31, 76, 82, 83*). Our present study suggests that HOP/Sti1 has a post-maturation role in case of CRAF kinase as evident from unaltered Ser-621 phosphorylation level (Fig.1Aii), a maturation marker of CRAF kinase (*38*), however, a significant reduction of CRAF kinase activity was observed under Sti1/HOP depleted condition (Fig. 1Ai and D). Moreover, HOP depletion does not affect CRAF kinase stability as evident from the cycloheximide chase analysis (Fig. S7). On contrary, impairment in Cdc37 (another client recruiter co-chaperone of Hsp90) functioning affects CRAF maturation, steady-state expression level and its kinase activity (*17*) suggesting a differential role of both co-chaperones HOP and Cdc37 towards kinase processing. HOP is an integral component of CRAF complex responsible for CRAF kinase activity which cannot be supplemented by Cdc37 (Fig 6C and E). Since both co-chaperones HOP and Cdc37 act as recruiter of Hsp90 to its client, hence, depletion of HOP reduces Hsp90 binding to CRAF (Fig. 6A) as that of Cdc37 (*17*). However, unaltered pS621 level of CRAF kinase in absence of HOP may be due to the presence of Cdc37 or by the joint action of Hsp70-Hsp90 which can supplement the role of HOP during folding as reported recently (*84*). Our data shows that HOP recruits Hsp90 to CRAF during activation of MAPK pathway (Fig. 5E), therefore, HOP silencing impairs CRAF signaling (Fig. 1F) which is not rescued by Cdc37 (Fig 6E). This observation strongly suggests that recruitment of Hsp90 during CRAF activation is facilitated by HOP, while the initial/continuous association of Hsp90 to CRAF complex is driven by both Cdc37 and HOP. Collectively, the differential behaviors of two co-chaperones towards CRAF-Hsp90 assemblage strengthen the fact of two independent recruitment events of Hsp90 to a single client CRAF kinase. However, further molecular evidence is required to identify the exact mechanism behind the dual recruitments of Hsp90 to its client CRAF kinase.

The precise functioning of HOP in client activation is guided by its different structural domains (*54*). Sti1 fragment containing the TPR2A-2B domain is solely unable to keep the client in its active state, although the indicated domain (TPR2A-2B) still interacts with both Hsp70 and Hsp90 (*76*). Importantly, the presence of intact TPR2A-2B-DP2 domain in Sti1 fragment is sufficient for complete activation of Glucocorticoid receptor (GR) depicting a critical role of DP2 domain in HOP-dependent client activation (*41*). Cryo-EM study confirms the interaction of DP2 domain of HOP with GR, and both protomers of Hsp90, thereby, maintaining Hsp90 in semi-closed conformation in client loading complex (*76*). Our data redefines that the integrity of TPR2A-2B-DP2 domain of HOP is necessary and sufficient for CRAF activation enabling Hsp90 recruitment to the CRAF complex and thereby regulating its activation (Figs. 3F and 4Bii). Notably, HOP^WT^ was unable to rescue the intrinsic CRAF activity in comparison to exogenously expressed Sti1 in *sti1Δ* background (Fig. 3E). The probable reason behind such observation was may be due to differences in functional efficiency in yeast system between exogenously expressed HOP and Sti1. Our further findings confirm that HOP is essential to keep the activation competent CRAF complex by recruiting both Hsp90 and actin during MAPK activation (Fig. 4D). Therefore, our study not only redefines the essential requirement of co-chaperone like HOP/Sti1 in CRAF kinase activation, but also uncovers the mechanistic insight for why HOP is essential during CRAF signaling. During this study, our experimental data opens two possible mechanisms behind the continuous association of HOP in the CRAF complex. Being an adaptor co-chaperone, HOP might be involved to accelerate the rate of Hsp90 recruitment to the CRAF complex during signaling. Alternatively, being a component in CRAF complex HOP/Sti1 might keep cytosolic CRAF in such a conformation regulating its basal kinase activity, for which initial Hsp90 attachment is not sufficient to keep the kinase in its active conformation. However, HOP-mediated increased recruitment of Hsp90 to CRAF complex also unravels an existence of parallel possibility that HOP might engage another Hsp90 molecule to CRAF complex, although this observation requires further investigation for in detailed mechanism.

HOP interacts with C-terminal EEVD domain of both Hsp70 and Hsp90 via its TPR motif and involves both in protein folding and degradation (*85, 86*). Recently, Hsp70-HOP-Hsp90 ternary complex is found for optimal assembly and maintenance of 26S/30S proteasome, which switches the proteostatic balance from folding to degradation (*84*). Since Hsp70-HOP interaction may target client protein for degradation we checked whether HOP has any role in degradation of CRAF kinase. To explore that geldanamycin (GA) induced degradation of CRAF was monitored in HEK293T cells upon downregulation of HOP by siRNA. However, unaltered CRAF level in HOP silenced condition compared to control suggests that HOP does not involve in CRAF degradation defining its functional role in maintaining CRAF activation (Fig. S7).

Although co-chaperone mediated *in-vitro* client complex formation is well documented in several earlier studies, while co-chaperone and its client post-maturation events need careful attention. This study establishes that a single co-chaperone HOP can regulate kinase functioning differently at two distinct layers of client chaperoning process. First, during the kinase client transfer in folding and maturation step, and second, at the time of kinase’s participation in signaling process. Here, we identify that HOP/Sti1 acts as a post-maturation Hsp90 recruiter protein that interacts with both Hsp90 and CRAF kinase to produce an effective transport complex during MAPK signaling. In addition, HOP/Sti1 is also essential to keep cytosolic CRAF in an activation-competent conformation (Fig. 7). Overall, our study demonstrates HOP as a novel CRAF kinase regulator, which specifically monitors Hsp90-dependent RAF kinase activation and establishes the physiological importance of HOP-Hsp90 interaction with kinase beyond its role in client folding and maturation. Moreover, dependence of mutant CRAF functioning on HOP opens an alternative target to mitigate the CRAF mediated Rasopathies.

**FIGURE 7.**
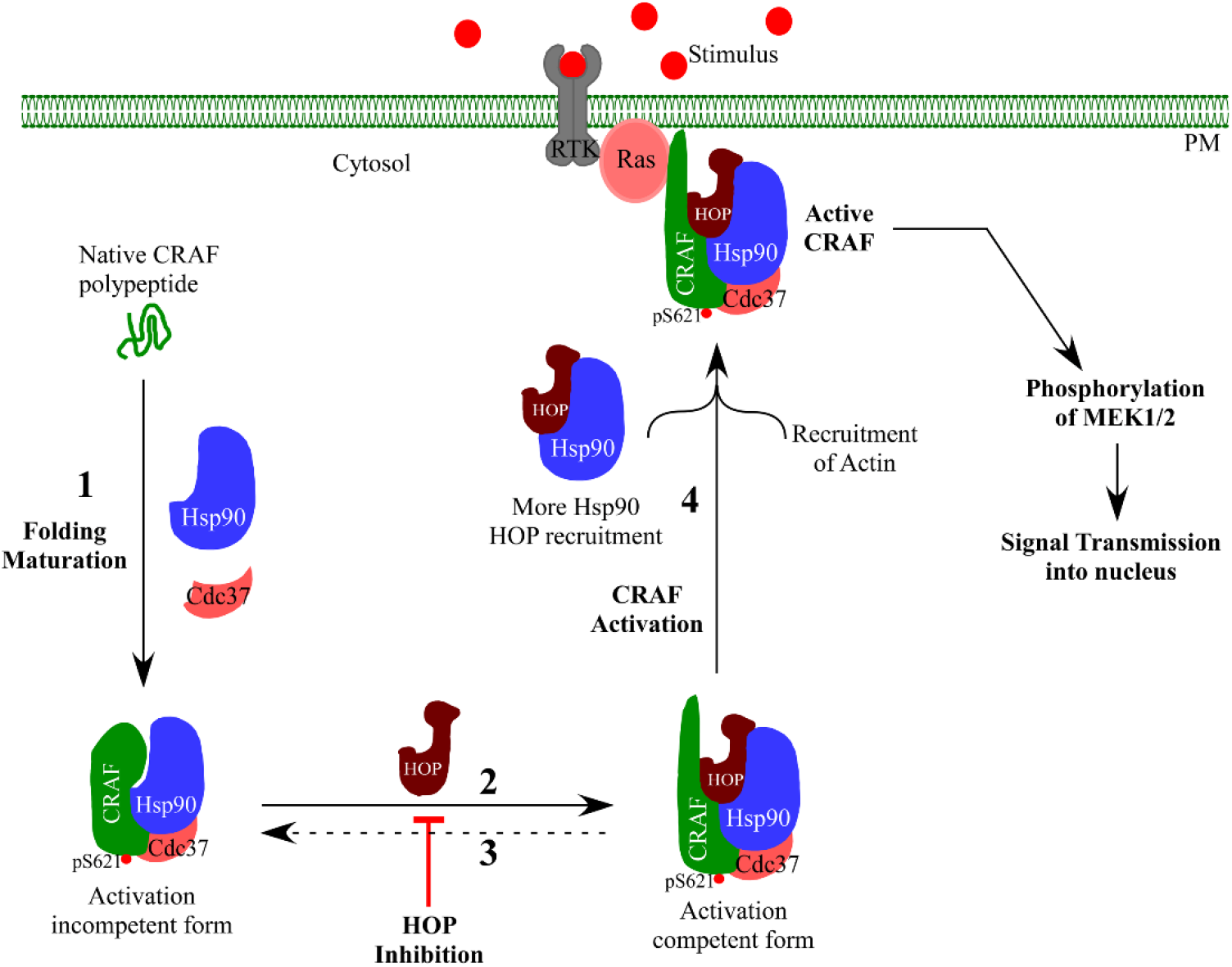
The proposed model describing the role of HOP in CRAF activation: The nascent CRAF polypeptide is folded and matured by Hsp90-Cdc37 chaperone machinery. Here, Cdc37 recruits Hsp90 to CRAF and facilitates its autophosphorylation at Ser-621 residue. Eventually, the newly synthesized kinase molecule achieves activation incompetent state (1). Due to post-folding entry of HOP into CRAF-Hsp90 complex, the kinase attains its activation competent state (2). However, depletion/ inhibition of HOP alters CRAF basal activity keeping CRAF-Hsp90 interaction intact. Therefore, HOP silencing facilitates the abundance of activation incompetent CRAF form, as shown by the reverse dotted line in the model (3). During MAPK signaling (presence of stimulus), a simultaneous recruitment of Hsp90, HOP and actin occurs to produce effective active CRAF transport signalosome to transmit growth signal inside the nucleus (4). Notably, at this step HOP keeps Hsp90 in an open conformation to facilitate its recruitment inside the CRAF complex suggesting how co-chaperone HOP assists Hsp90 during client kinase activation. Inhibition of HOP thus restricts MAPK activation by compromising Hsp90-CRAF association. Here, RTK and PM represent receptor tyrosine kinase and plasma membrane.

## Materials and Methods

### Yeast strains, media and chemicals and antibodies

BY4741 (MATa his3Δ0 leu2Δ0 met15Δ0 ura3Δ0) yeast strain was used in this study. Hsp90 co-chaperone knock out yeast strains, *aha1∷KanMx, sti1∷KanMx* and *sba1∷KanMx* were obtained from the yeast knock out library. Yeast cells were grown in either YPD (1% Bacto-yeast extract, 2% Bacto-peptone, 2% dextrose) media or in selective dropout media. Yeast transformation was carried out by using standard Lithium acetate protocol. G418 sulfate was obtained from Sigma (St. Louis, MO) and used at a final concentration of 400 μg /ml. Antibodies were obtained from the following sources: anti-FLAG 1:2500 (Cell Signaling Technology (CST) #14793, Sigma #F1804, #F7425), anti-HA, 1:5000 (Sigma #H3663); anti-MYC, 1:5000 (CST #2276); anti-HOP, 1:5000 (CST #5670); anti-MEK1/2, 1:5000 (CST #4694); anti-pMEK1/2, 1:5000 (CST #9154); anti-EGFR, anti-ERBB2, 1:1000 (Santa Cruz Biotechnology #SC373746, CST #2165S) and their phospho antibodies, 1:1000 (Santa cruz #SC23420R, CST #2243S); anti-actin, 1:5000 (CST #D18C11); anti-CRAF and anti-BRAF (BD Bioscience #610152 and #612375); anti-GAPDH (Biobharati Life Sciences #BB-AB0060); anti-Ras (Thermo Fisher scientific #MA1-012). Protein A sepharose bead was purchased from GE healthcare. On target plus smart pool HOP siRNA (# L-019802-00-0005) and on target plus non-targeting pool siRNA (#D-001810-10-05) were purchased from Dharmacon.

### Cell lines, Cell culture media and Transfection

HEK293T cell was obtained from National Centre for Cell Science Cell Repository (Pune, India) and maintained in a 5% CO_2_ containing humified 37°C incubator in Dulbecco’s modified Eagles medium (DMEM, Himedia, #AL007G-500ML) supplemented with 10% FBS (Gibco #16000044), 150μg/ml penstrep, 250μg/ml amphotericin B and 50 mg/ml of gentamicin. For transient transfection, plasmids were generally transfected at 50-60% of confluency using PEI (polyethyleneimine, Polysciences Inc #23966-2)/ Lipofectamine LTX (Invitrogen #15338100) according to the manufacturer’s protocol. Lipofectamine 2000 (Invitrogen #11668019) transfection reagent was used for siRNA mediated gene silencing.

### Plasmids

Follow Supporting Information Table S1.

### Western blotting and Immunoprecipitation

The expression of CRAF and HOP proteins in mammalian cell and yeast was assayed by lysing the cells using RIPA cell lysis buffer (20 mM Tris, pH-7.5, 150 mM NaCl, and 0.l % NP-40, 1%SDS along with 10mM PMSF, 1X phosphatase inhibitor ((Thermo Pierce #1862495)) and 1X protease inhibitor cocktail (Roche #04693116001). Cell lysates were clarified by centrifugation at 13,000 rpm for 15 mins at 4^0^C. The protein concentrations in cell lysates were measured by BCA method. For immunoprecipitation, cell lysates (50mM Tris, pH-8.0, 150mM NaCl, 0.2% Triton X, 10mM PMSF, 1X phosphatase inhibitor and 1X protease inhibitor) were incubated with antibody for 16 hrs at 4^0^C. Protein-A sepharose bead was added and again incubated for 2 hrs at 4^0^C. The beads were then washed with buffer (50mM Tris, pH-8.0, 150mM NaCl, 0.2% Triton X, 10mM PMSF) and resuspended in 2x SDS sample loading buffer. The immune-complex was separated onto 10% SDS PAGE and analyzed by western blotting.

### CRAF Kinase assay

CRAF kinase activity was measured as described previously (*17*) and according to the manufacture’s protocol (Millipore). Briefly, Flag-CRAF/Endogenous CRAF was immunoprecipitated by anti-Flag/anti-CRAF antibody from yeast (Flag-CRAF) or mammalian cell lysates (Flag-CRAF/Endogenous CRAF). The immune-complex was pulled down by protein-A sepharose beads by incubating at 4°C for 2 hrs. The beads were then washed three times with buffer ( 50mM Tris, pH-8.0, 150mM NaCl, 0.2% Triton X, 10mM PMSF) and kinase assay buffer ( 20mM MOPS pH7.2, 25 mM β–Glycerophosphate, 5mM EGTA, 1mM Sodium ortho-vanadate, 1mM Dithiothreitol) was added along with Mg^+2^ATP. Purified inactive MEK1 was added as a kinase substrate and incubated at 30°C in a water bath with gentle shaking for 1 hr. The activity of immunoprecipitated CRAF protein was assessed by the level of MEK phosphorylation (pMEK1) in western blot.

### Detection of extracellular HOP secretion

To detect whether HOP is secreted out from the cells upon over-expression, semi confluent HEK293T cells were transfected with Myc-HOP plasmid. After transfection, cells are cultured in serum-free media for 24h and 48h respectively. After this period, conditioned media was isolated and centrifuged at 3000g. The supernatant was collected and lyophilized. The centrifuged cell pellet and the lyophilized fraction were dissolved in 2X sample buffer and analyzed by western blot.

### Immunofluoroscence

Immunofluoresence experiments were carried out by transfecting the cells transiently with plasmids at 60-70% of confluence for 12 hrs. The transfected cells were then seeded onto poly-L-lysine coated sterile cover slips and kept for another 36 hours. The cells on cover slips were washed three times with 1X PBS and fixed with 3.7% formaldehyde. The cells were then permeabilized with 1% triton-X and blocked with 5% BSA for an hour at RT. The cover slips were washed twice with 1X PBS and probed with primary antibody at 4°C for overnight. Next day, cover slips were washed with 1X PBS and probed with secondary antibody for 1 hr at RT. After washing three times with 1X PBS, cover slips were then mounted onto glass slide with vectashield (Invitrogen) and examined under confocal microscope.

### Statistical analysis

All the statistical analyses were performed using GraphPad Prism. Two tailed student t-test or one-way anova with Tukey’s Multiple Comparison Test was used for analyzing the significance. Data was expressed as mean ± SD. For all tests, a p value less than 0.05 was considered as statistically significant.

## Acknowledgements

We are grateful to Professor Johannes Buchner (Technische Universitat Munchen, Munich, Germany) for providing mammalian HOP plasmid and yeast Sti1 constructs. This work was supported by Department of Biotechnology, Government of India, (BT/PR6544/BRB/10/1151/2012 to AKM) and Bose Institute Intramural funding. The author thanks to Council of Scientific & Industrial Research (CSIR) for fellowship to NG.

## Author contribution

SM and AKM conceived the idea; NG, SM, AKM designed the experiments, NG performed most of the experiments; SM, SR contributed in some part of experiments; NG, SM, and AKM analyzed the results; SM, NG, AKM wrote the manuscript. All the authors reviewed and corrected the manuscript.

## Abbreviations

CRAF: Raf1 kinase
HOP: Hsp70/Hsp90 organizing protein;
Hsp: heat shock protein;
MAPK: mitogen-activated protein kinase;
EGF: epidermal growth factor;
EGFR: EGF receptor;
GFP: green fluorescent protein;
DsRed: red fluorescent protein;
GPD: glyceraldehyde-3-phosphate dehydrogenase;
Co-IP: co-immunoprecipitation;

## References

1. A. Plotnikov, E. Zehorai, S. Procaccia, R. Seger, The MAPK cascades: signaling components, nuclear roles and mechanisms of nuclear translocation. Biochim Biophys Acta 1813, 1619–1633 (2010).

2. S. M. Storm, J. L. Cleveland, U. R. Rapp, Expression of raf family proto-oncogenes in normal mouse tissues. Oncogene 5, 345–351 (1990).

3. M. Huser et al., MEK kinase activity is not necessary for Raf-1 function. EMBO J 20, 1940–1951 (2001).

4. L. Wojnowski et al., Craf-1 protein kinase is essential for mouse development. Mech Dev 76, 141–149 (1998).

5. M. Zenker, Clinical overview on RASopathies. Am J Med Genet C Semin Med Genet, (2022).

6. A. Noeparast et al., CRAF mutations in lung cancer can be oncogenic and predict sensitivity to combined type II RAF and MEK inhibition. Oncogene 38, 5933–5941 (2019).

7. H. Chong, H. G. Vikis, K. L. Guan, Mechanisms of regulating the Raf kinase family. Cell Signal 15, 463–469 (2003).

8. A. S. Dhillon, W. Kolch, Untying the regulation of the Raf-1 kinase. Arch Biochem Biophys 404, 3–9 (2002).

9. N. Dumaz, R. Marais, Protein kinase A blocks Raf-1 activity by stimulating 14-3-3 binding and blocking Raf-1 interaction with Ras. J Biol Chem 278, 29819–29823 (2003).

10. M. Molzan et al., Impaired binding of 14-3-3 to C-RAF in Noonan syndrome suggests new approaches in diseases with increased Ras signaling. Mol Cell Biol 30, 4698–4711 (2010).

11. G. Tzivion, Z. Luo, J. Avruch, A dimeric 14-3-3 protein is an essential cofactor for Raf kinase activity. Nature 394, 88–92 (1998).

12. D. Abraham et al., Raf-1-associated protein phosphatase 2A as a positive regulator of kinase activation. J Biol Chem 275, 22300–22304 (2000).

13. N. Grammatikakis, J. H. Lin, A. Grammatikakis, P. N. Tsichlis, B. H. Cochran, p50(cdc37) acting in concert with Hsp90 is required for Raf-1 function. Mol Cell Biol 19, 1661–1672 (1999).

14. J. L. Holmes, S. Y. Sharp, S. Hobbs, P. Workman, Silencing of HSP90 cochaperone AHA1 expression decreases client protein activation and increases cellular sensitivity to the HSP90 inhibitor 17-allylamino-17-demethoxygeldanamycin. Cancer Res 68, 1188–1197 (2008).

15. O. M. Grbovic et al., V600E B-Raf requires the Hsp90 chaperone for stability and is degraded in response to Hsp90 inhibitors. Proc Natl Acad Sci U S A 103, 57–62 (2006).

16. T. W. Schulte, M. V. Blagosklonny, C. Ingui, L. Neckers, Disruption of the Raf-1-Hsp90 molecular complex results in destabilization of Raf-1 and loss of Raf-1-Ras association. J Biol Chem 270, 24585–24588 (1995).

17. S. Mitra, B. Ghosh, N. Gayen, J. Roy, A. K. Mandal, Bipartite Role of Heat Shock Protein 90 (Hsp90) Keeps CRAF Kinase Poised for Activation. J Biol Chem 291, 24579–24593 (2016).

18. L. Muller, A. Schaupp, D. Walerych, H. Wegele, J. Buchner, Hsp90 regulates the activity of wild type p53 under physiological and elevated temperatures. J Biol Chem 279, 48846–48854 (2004).

19. D. Picard, Chaperoning steroid hormone action. Trends Endocrinol Metab 17, 229–235 (2006).

20. W. B. Pratt, D. O. Toft, Regulation of signaling protein function and trafficking by the hsp90/hsp70-based chaperone machinery. Exp Biol Med (Maywood) 228, 111–133 (2003).

21. M. Taipale et al., Quantitative analysis of HSP90-client interactions reveals principles of substrate recognition. Cell 150, 987–1001 (2012).

22. T. Scheibel, J. Buchner, The Hsp90 complex--a super-chaperone machine as a novel drug target. Biochem Pharmacol 56, 675–682 (1998).

23. J. Li, K. Richter, J. Reinstein, J. Buchner, Integration of the accelerator Aha1 in the Hsp90 co-chaperone cycle. Nat Struct Mol Biol 20, 326–331 (2013).

24. K. Richter, S. Walter, J. Buchner, The Co-chaperone Sba1 connects the ATPase reaction of Hsp90 to the progression of the chaperone cycle. J Mol Biol 342, 1403–1413 (2004).

25. S. M. Roe et al., The Mechanism of Hsp90 regulation by the protein kinase-specific cochaperone p50(cdc37). Cell 116, 87–98 (2004).

26. D. F. Smith et al., Identification of a 60-kilodalton stress-related protein, p60, which interacts with hsp90 and hsp70. Mol Cell Biol 13, 869–876 (1993).

27. A. J. Caplan, A. K. Mandal, M. A. Theodoraki, Molecular chaperones and protein kinase quality control. Trends Cell Biol 17, 87–92 (2007).

28. L. M. Karnitz, S. J. Felts, Cdc37 regulation of the kinome: when to hold ‘em and when to fold’em. Sci STKE 2007, pe22 (2007).

29. H. C. Chang, D. F. Nathan, S. Lindquist, In vivo analysis of the Hsp90 cochaperone Sti1 (p60). Mol Cell Biol 17, 318–325 (1997).

30. P. E. Carrigan, D. L. Riggs, M. Chinkers, D. F. Smith, Functional comparison of human and Drosophila Hop reveals novel role in steroid receptor maturation. J Biol Chem 280, 8906–8911 (2005).

31. P. Lee, A. Shabbir, C. Cardozo, A. J. Caplan, Sti1 and Cdc37 can stabilize Hsp90 in chaperone complexes with a protein kinase. Mol Biol Cell 15, 1785–1792 (2004).

32. Y. Song, D. C. Masison, Independent regulation of Hsp70 and Hsp90 chaperones by Hsp70/Hsp90-organizing protein Sti1 (Hop1). J Biol Chem 280, 34178–34185 (2005).

33. B. D. Johnson, R. J. Schumacher, E. D. Ross, D. O. Toft, Hop modulates Hsp70/Hsp90 interactions in protein folding. J Biol Chem 273, 3679–3686 (1998).

34. H. Kubota et al., Increased expression of co-chaperone HOP with HSP90 and HSC70 and complex formation in human colonic carcinoma. Cell Stress Chaperones 15, 1003–1011 (2010).

35. N. Walsh et al., RNAi knockdown of Hop (Hsp70/Hsp90 organising protein) decreases invasion via MMP-2 down regulation. Cancer Lett 306, 180–189 (2011).

36. T. H. Wang et al., Stress-induced phosphoprotein 1 as a secreted biomarker for human ovarian cancer promotes cancer cell proliferation. Mol Cell Proteomics 9, 1873–1884 (2010).

37. J. D. Sims, J. McCready, D. G. Jay, Extracellular heat shock protein (Hsp)70 and Hsp90alpha assist in matrix metalloproteinase-2 activation and breast cancer cell migration and invasion. PLoS One 6, e18848 (2011).

38. C. Noble et al., CRAF autophosphorylation of serine 621 is required to prevent its proteasome-mediated degradation. Mol Cell 31, 862–872 (2008).

39. A. Wolmarans, B. Lee, L. Spyracopoulos, P. LaPointe, The Mechanism of Hsp90 ATPase Stimulation by Aha1. Sci Rep 6, 33179 (2016).

40. Y. Fang, A. E. Fliss, J. Rao, A. J. Caplan, SBA1 encodes a yeast hsp90 cochaperone that is homologous to vertebrate p23 proteins. Mol Cell Biol 18, 3727–3734 (1998).

41. A. Rohl et al., Hop/Sti1 phosphorylation inhibits its co-chaperone function. EMBO Rep 16, 240–249 (2015).

42. T. A. Americo, L. B. Chiarini, R. Linden, Signaling induced by hop/STI-1 depends on endocytosis. Biochem Biophys Res Commun 358, 620–625 (2007).

43. B. K. Eustace, D. G. Jay, Extracellular roles for the molecular chaperone, hsp90. Cell Cycle 3, 1098–1100 (2004).

44. J. Avruch, X. F. Zhang, J. M. Kyriakis, Raf meets Ras: completing the framework of a signal transduction pathway. Trends Biochem Sci 19, 279–283 (1994).

45. A. Kauffmann-Zeh et al., Suppression of c-Myc-induced apoptosis by Ras signalling through PI(3)K and PKB. Nature 385, 544–548 (1997).

46. M. F. Favata et al., Identification of a novel inhibitor of mitogen-activated protein kinase kinase. J Biol Chem 273, 18623–18632 (1998).

47. L. Adnane, P. A. Trail, I. Taylor, S. M. Wilhelm, Sorafenib (BAY 43-9006, Nexavar), a dual-action inhibitor that targets RAF/MEK/ERK pathway in tumor cells and tyrosine kinases VEGFR/PDGFR in tumor vasculature. Methods Enzymol 407, 597–612 (2006).

48. S. J. Leevers, H. F. Paterson, C. J. Marshall, Requirement for Ras in Raf activation is overcome by targeting Raf to the plasma membrane. Nature 369, 411–414 (1994).

49. C. K. Weber, J. R. Slupsky, H. A. Kalmes, U. R. Rapp, Active Ras induces heterodimerization of cRaf and BRaf. Cancer Res 61, 3595–3598 (2001).

50. P. E. Carrigan et al., Multiple domains of the co-chaperone Hop are important for Hsp70 binding. J Biol Chem 279, 16185–16193 (2004).

51. S. Daniel et al., Nuclear translocation of the phosphoprotein Hop (Hsp70/Hsp90 organizing protein) occurs under heat shock, and its proposed nuclear localization signal is involved in Hsp90 binding. Biochim Biophys Acta 1783, 1003–1014 (2008).

52. K. Bhattacharya, L. Bernasconi, D. Picard, Luminescence resonance energy transfer between genetically encoded donor and acceptor for protein-protein interaction studies in the molecular chaperone HSP70/HSP90 complexes. Sci Rep 8, 2801 (2018).

53. M. Castelli et al., Phosphorylation of the Hsp90 Co-Chaperone Hop Changes its Conformational Dynamics and Biological Function. J Mol Biol 435, 167931 (2022).

54. A. B. Schmid et al., The architecture of functional modules in the Hsp90 co-chaperone Sti1/Hop. EMBO J 31, 1506–1517 (2012).

55. D. R. Southworth, D. A. Agard, Client-loading conformation of the Hsp90 molecular chaperone revealed in the cryo-EM structure of the human Hsp90:Hop complex. Mol Cell 42, 771–781 (2011).

56. A. Rohl, J. Rohrberg, J. Buchner, The chaperone Hsp90: changing partners for demanding clients. Trends Biochem Sci 38, 253–262 (2013).

57. L. Stepanova et al., Induction of human Cdc37 in prostate cancer correlates with the ability of targeted Cdc37 expression to promote prostatic hyperplasia. Oncogene 19, 2186–2193 (2000).

58. L. Whitesell, S. L. Lindquist, HSP90 and the chaperoning of cancer. Nat Rev Cancer 5, 761–772 (2005).

59. L. S. Steelman et al., Roles of the Raf/MEK/ERK and PI3K/PTEN/Akt/mTOR pathways in controlling growth and sensitivity to therapy-implications for cancer and aging. Aging (Albany NY) 3, 192–222 (2011).

60. G. Hatzivassiliou et al., RAF inhibitors prime wild-type RAF to activate the MAPK pathway and enhance growth. Nature 464, 431–435 (2010).

61. P. I. Poulikakos, C. Zhang, G. Bollag, K. M. Shokat, N. Rosen, RAF inhibitors transactivate RAF dimers and ERK signalling in cells with wild-type BRAF. Nature 464, 427–430 (2010).

62. J. Hu et al., Allosteric activation of functionally asymmetric RAF kinase dimers. Cell 154, 1036–1046 (2013).

63. M. J. Garnett, S. Rana, H. Paterson, D. Barford, R. Marais, Wild-type and mutant B-RAF activate C-RAF through distinct mechanisms involving heterodimerization. Mol Cell 20, 963–969 (2005).

64. P. T. Wan et al., Mechanism of activation of the RAF-ERK signaling pathway by oncogenic mutations of B-RAF. Cell 116, 855–867 (2004).

65. M. Sanclemente et al., c-RAF Ablation Induces Regression of Advanced Kras/Trp53 Mutant Lung Adenocarcinomas by a Mechanism Independent of MAPK Signaling. Cancer Cell 33, 217–228 e214 (2018).

66. M. T. Blasco et al., Complete Regression of Advanced Pancreatic Ductal Adenocarcinomas upon Combined Inhibition of EGFR and C-RAF. Cancer Cell 35, 573–587 e576 (2019).

67. A. Venkatanarayan et al., CRAF dimerization with ARAF regulates KRAS-driven tumor growth. Cell Rep 38, 110351 (2022).

68. X. Wu et al., Increased BRAF heterodimerization is the common pathogenic mechanism for noonan syndrome-associated RAF1 mutants. Mol Cell Biol 32, 3872–3890 (2012).

69. N. R. Michaud et al., KSR stimulates Raf-1 activity in a kinase-independent manner. Proc Natl Acad Sci U S A 94, 12792–12796 (1997).

70. D. K. Morrison, R. E. Cutler, The complexity of Raf-1 regulation. Curr Opin Cell Biol 9, 174–179 (1997).

71. M. Wartmann, R. J. Davis, The native structure of the activated Raf protein kinase is a membrane-bound multi-subunit complex. J Biol Chem 269, 6695–6701 (1994).

72. Y. Morishima et al., The Hsp organizer protein hop enhances the rate of but is not essential for glucocorticoid receptor folding by the multiprotein Hsp90-based chaperone system. J Biol Chem 275, 6894–6900 (2000).

73. Y. J. Tao, W. Zheng, Chaperones and the maturation of steroid hormone receptor complexes. Oncotarget 2, 104–106 (2011).

74. P. Sahasrabudhe, J. Rohrberg, M. M. Biebl, D. A. Rutz, J. Buchner, The Plasticity of the Hsp90 Co-chaperone System. Mol Cell 67, 947–961 e945 (2017).

75. C. M. Noddings, R. Y. Wang, J. L. Johnson, D. A. Agard, Structure of Hsp90-p23-GR reveals the Hsp90 client-remodelling mechanism. Nature 601, 465–469 (2022).

76. R. Y. Wang et al., Structure of Hsp90-Hsp70-Hop-GR reveals the Hsp90 client-loading mechanism. Nature 601, 460–464 (2022).

77. S. J. Arlander et al., Chaperoning checkpoint kinase 1 (Chk1), an Hsp90 client, with purified chaperones. J Biol Chem 281, 2989–2998 (2006).

78. Z. Chen et al., Autocrine STIP1 signaling promotes tumor growth and is associated with disease outcome in hepatocellular carcinoma. Biochem Biophys Res Commun 493, 365–372 (2017).

79. V. M. Longshaw, M. Baxter, M. Prewitz, G. L. Blatch, Knockdown of the co-chaperone Hop promotes extranuclear accumulation of Stat3 in mouse embryonic stem cells. Eur J Cell Biol 88, 153–166 (2009).

80. K. Naruse, E. Matsuura-Suzuki, M. Watanabe, S. Iwasaki, Y. Tomari, In vitro reconstitution of chaperone-mediated human RISC assembly. RNA 24, 6–11 (2018).

81. S. Alvira et al., Structural characterization of the substrate transfer mechanism in Hsp70/Hsp90 folding machinery mediated by Hop. Nat Commun 5, 5484 (2014).

82. F. H. Schopf, M. M. Biebl, J. Buchner, The HSP90 chaperone machinery. Nat Rev Mol Cell Biol 18, 345–360 (2017).

83. E. E. Boczek et al., Conformational processing of oncogenic v-Src kinase by the molecular chaperone Hsp90. Proc Natl Acad Sci U S A 112, E3189–3198 (2015).

84. K. Bhattacharya et al., The Hsp70-Hsp90 co-chaperone Hop/Stip1 shifts the proteostatic balance from folding towards degradation. Nat Commun 11, 5975 (2020).

85. A. B. Abildgaard et al., Co-Chaperones in Targeting and Delivery of Misfolded Proteins to the 26S Proteasome. Biomolecules 10, (2020).

86. F. Eisele et al., An Hsp90 co-chaperone links protein folding and degradation and is part of a conserved protein quality control. Cell Rep 35, 109328 (2021).

