## Supplemental Data for "Hsp70/Hsp90 organizing protein (HOP) maintains CRAF kinase activity and regulates MAPK signaling by enhancing Hsp90-CRAF association"

### Supplementary Information

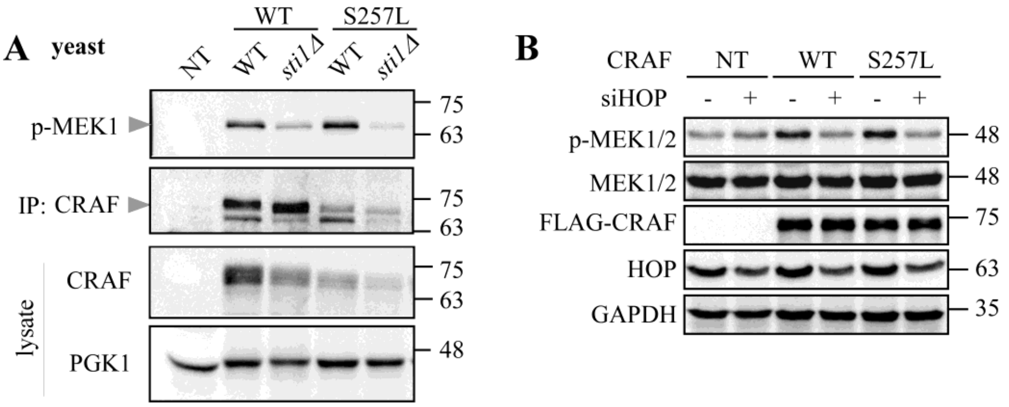


FIGURE S1. **Activity of hyperactive CRAF mutant is reduced upon HOP depletion. (A)** Overexpressed WT and mutant CRAF(S257L) from WT and *sti1Δ* yeast cells were pulled using an anti-CRAF antibody. The kinase activity of immunoprecipited CRAF was analyzed using *in-vitro* CRAF kinase assay and measured by the intensity of pMEK1 to the intensity of CRAF pulled. **(B)** HOP silenced and non-silenced HEK293T cells were transiently transfected with Flag-tagged WT and mutant CRAF (S257L). Cells were lysed to test the activity MAPK pathway by the magnitude of the pMEK1/2 band to the tMEK1/2 band. Flag-CRAF and HOP represent the amount of expressed proteins respectively. GAPDH served as the loading control.

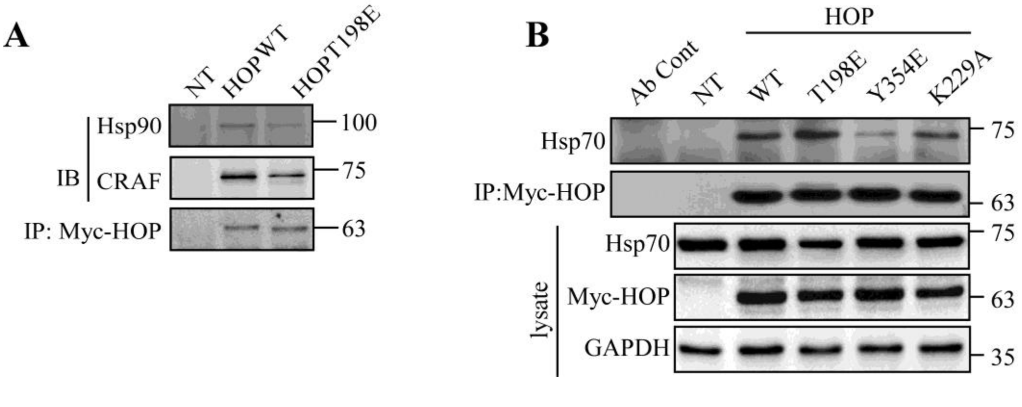


FIGURE S2. **HOP mutants, T198E and Y354E cannot interact with Hsp90 and Hsp70 respectively**. Myc-HOP and its mutants, T198E, Y354E and K229A respectively were transformed in HEK293T cells. Myc-HOP was immunoprecipitated and the binding of Hsp90 (**A**), Hsp70 (**B**) and CRAF was evaluated with the respective antibodies. NT represents non-transfected cells.

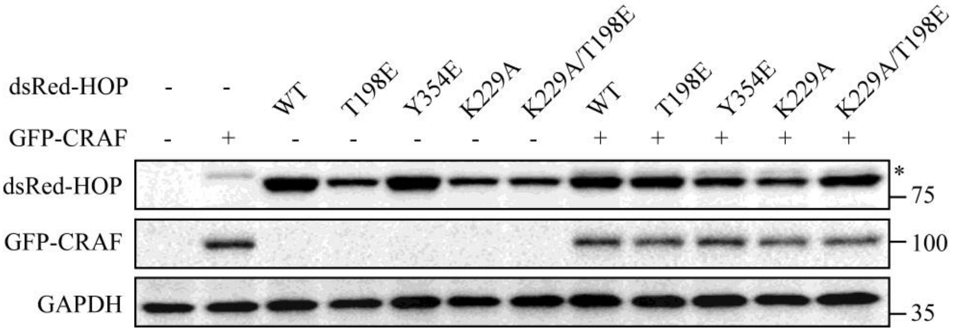


FIGURE S3. Western blot analysis of HEK293T cell lysates shows the expression of GFP-CRAF and respective DsRed-HOP mutants as described in Fig. 3D.

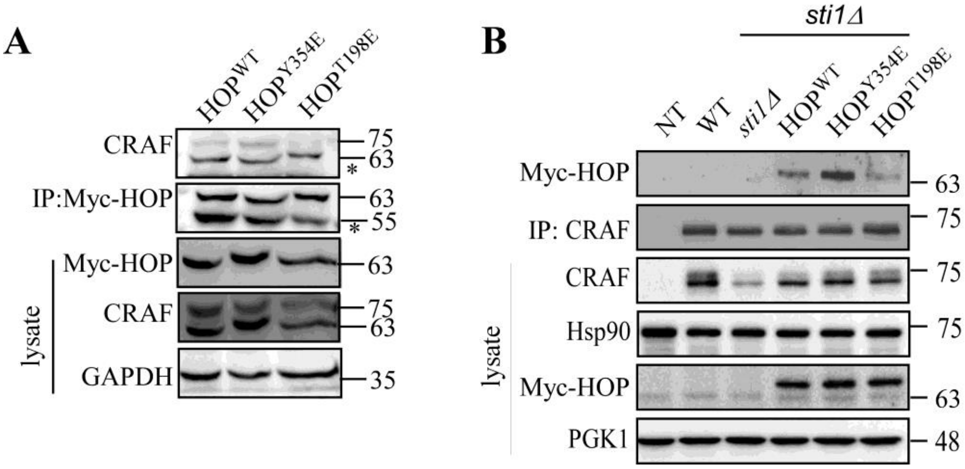


FIGURE S4. **HOP$^{T198E}$ mutant showed reduced interaction with CRAF**. Indicated Myc-HOP constructs were over-expressed in HEK293T cells (A) and also in *sti1Δ* yeast cells bearing Flag-CRAF (B). Either Myc-HOP or Flag-CRAF was immunoprecipitated and the association of endogenous CRAF or Myc-HOP was detected by western blotting with anti-CRAF and anti-myc antibodies respectively. Lysate showed the expression of the respective protein. * Indicates the nonspecific protein.

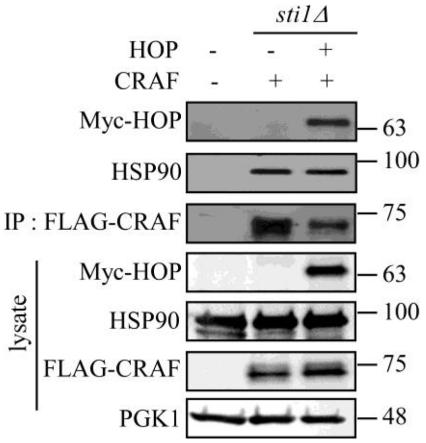


FIGURE S5. **HOP remains in complex with CRAF**. Flag-CRAF was expressed in *sti1Δ* yeast cells in absence and presence of Myc-HOP. CRAF was immunoprecipitated with anti-Flag antibody and its association with HOP and Hsp90 were observed by western blotting using anti-Myc and anti-Hsp90 antibodies respectively. Lysate shows the expression of indicated proteins, PGK1 was used as loading control.

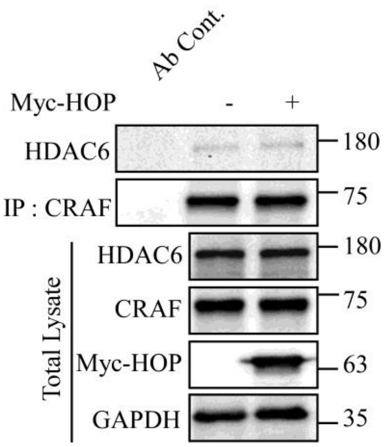


FIGURE S6. **HDAC6 is not involved in CRAF translocation**. Myc-HOP was expressed in HEK293T cells. Endogenous CRAF was immunoprecipitated with anti-CRAF antibody and the association of HDAC6 was checked by western blot analysis. Ab Cont. represents antibody control.

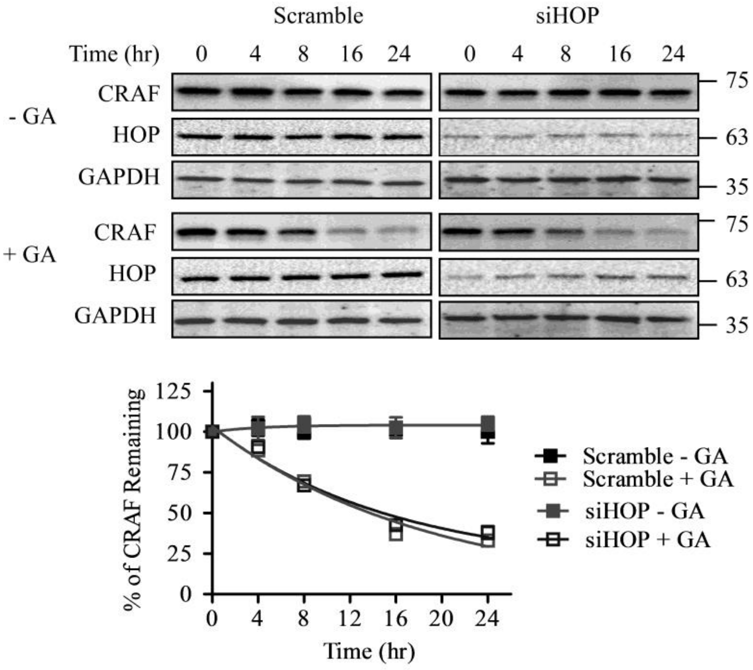


FIGURE S7. **HOP does not involve in CRAF degradation**. HOP silencing was carried out in HEK293T cells. Geldanamycin (GA) was added to the cells and the cells were taken at the indicated time. CRAF level was checked by western blot analysis with anti-CRAF antibody. CRAF level was quantified, normalized with GAPDH, and plotted in the graph. Error bar represents SD of two independent experiments.

Table S1. Plasmids used in the study

| Plasmids | Description | Reference |
|---|---|---|
| p425GPD Sti1<br>p425GPD HOP | Wild type Sti1, Wild type HOP, Leu 2-based expression plasmid, GPD promoter | Kind gift from Prof. Johannes Buchner , Technische Universität München, Munich, Germany. |
| pcDNA3 Flag-CRAF | Mammalian CRAF with N-terminal Flag-tag | Kind gift from Prof. Bruce Gelb, Mount Sinai School of Medicine, NY, USA. |
| p416GPD Flag-CRAF | Mammalian CRAF with N-terminal Flag tag, Ura 3-based expression plasmid | Mitra et al., 2016) |
| p413GPD Flag-CRAF | CRAF with N-terminal Flag tag, His 3-based expression plasmid | (Mitra et al., 2016) |
| p416GPD Flag-BRAF | Mammalian BRAF with N-terminal Flag tag | This study |
| p416GPD Myc-HOP | Mammalian wild type HOP with N-terminal Myc tag, GPD promoter (HindIII+XhoI) | This study |
| p416GPD Myc-HOP $^{T198E}$ | T198E mutated HOP with N-terminal Myc tag | This study |
| p416GPD Myc-HOP $^{Y354E}$ | Y354E mutated HOP with N-terminal Myc tag | This study |
| p416GPD Myc-HOP $^{\Delta DP2}$ | DP2 domain deleted HOP with N-terminal Myc tag, (HindIII+XhoI) | This study |
| p416GPD Myc-HOP $^{TPR2A\text{-}2B}$ | TPR 2A2B domain containing truncated HOP with N-terminal Myc tag, (HindIII+XhoI) | This study |
| p416GPD Myc-HOP $^{TPR2B\text{-}DP2}$ | TPR 2BDP2 domain containing truncated HOP with N-terminal Myc tag, (HindIII+XhoI) | This study |
| p416GPD Myc-HOP $^{TPR2A\text{-}2B\text{-}DP2}$ | TPR 2A2BDP2 domain containing truncated HOP with N-terminal Myc tag, (HindIII+XhoI) | This study |
| p416GPD Myc-HOP $^{TPR2A\text{-}2B\text{-}DP2}$ $_{K229A}$ | K229A mutated TPR 2A-2B-DP2 domain containing truncated HOP with N-terminal Myc tag, (HindIII+XhoI) | This study |
| p416GPD Myc-HOP $^{TPR2A\text{-}2B\text{-}DP2}$ $_{T198E/K229A/Y354E}$ | T198E/K229A/Y354E mutated TPR 2A-2B-DP2 domain containing truncated HOP with N-terminal Myc tag, (HindIII+XhoI) | This study |
| pcDNA3.1 Myc-HOP | Mammalian HOP with N-terminal Myc tag, CMV promoter (HindIII+XhoI) | This study |
| pcDNA3.1 Myc-HOP $^{T198E}$ | T198E mutated HOP with N-terminal Myc tag | This study |
| pcDNA3.1 Myc-HOP $^{K229A}$ | K229A mutated HOP with N-terminal Myc tag | This study |
| pcDNA3.1 Myc-HOP $^{Y354E}$ | Y354E mutated HOP with N-terminal Myc tag | This study |
| pcDNA3.1 Myc-HOP $^{\Delta DP2}$ | DP2 domain deleted HOP with N-terminal Myc tag, CMV promoter. (HindIII+XhoI) | This study |
| pcDNA3.1 Myc-HOP $^{TPR2A\text{-}2B}$ | TPR2A-2B domain containing truncated HOP with N-terminal Myc tag, (HindIII+Xho1) | This study |
| pcDNA3.1 Myc-HOP $^{TPR2B\text{-}DP2}$ | TPR2B-DP2 domain containing truncated HOP with N-terminal Myc tag, (HindIII+XhoI) | This study |
| pcDNA3.1 Myc-HOP $^{TPR2A\text{-}2B\text{-}DP2}$ | TPR2A-2B-DP2 domain containing truncated HOP with N-terminal Myc tag, (HindIII+XhoI) | This study |
| pcDNA3.1 Myc-HOP $^{TPR2A\text{-}2B\text{-}DP2}$ $_{K229A}$ | K229A mutated TPR2A-2B-DP2 domain containing truncated HOP with N-terminal Myc tag, (HindIII+XhoI) | This study |
| pcDNA3.1 Myc-HOP $^{TPR2A\text{-}2B\text{-}DP2}$ $_{T198E/K229A}$ | T198E/K229A mutated TPR 2A-2B-DP2 domain containing truncated HOP with N-terminal Myc tag, (HindIII+XhoI) | This study |
| pcDNA3.1 Myc-HOP $^{TPR2A\text{-}2B\text{-}DP2}$ $_{T198E/K229A/Y354E}$ | T198E/K229A/Y354E mutated TPR 2A-2B-DP2 domain containing truncated HOP with | This study |

| | | |
|---|---|---|
| | N-terminal Myc tag, (HindIII+XhoI) | |
| pEGFPC1-CRAF | Mammalian CRAF with N-terminal GFP tag, CMV promoter. | (Mitra et al., 2016) |
| pcDNA3 GFP-HRasG12V | Mutant mammalian HRas G12V with N-terminal GFP tag. (BamHI+NotI) | This study |
| pcDNA3.1 Flag BRAF | Mammalian BRAF with N-terminal Flag tag. | Addgene (#40775) |
| pDsRedC1- $HOP^{WT}$ | Mammalian wild type HOP with N-terminal DsRed tag, CMV promoter (XhoI+EcoRI) | This study |
| pDsRedC1- $HOP^{T198E}$ | T198E mutated HOP with N-terminal DsRed tag, (XhoI+EcoRI) | This study |
| pDsRedC1- $HOP^{K229A}$ | K229A mutated HOP with N-terminal DsRed tag, (XhoI+EcoRI) | This study |
| pDsRedC1- $HOP^{Y354E}$ | Y354E mutated HOP with N-terminal DsRed tag, CMV promoter (XhoI+EcoRI) | This study |
| pDsRedC1- $HOP^{T198E/K229A}$ | T198E/K229A mutated HOP with N-terminal DsRed tag, (XhoI+EcoRI) | This study |
